# Endothelium-on-a-chip for permeability studies in diabetic and diabetic kidney disease milieu

**DOI:** 10.64898/2026.09.18.752765

**Authors:** Vika Telle, Pim de Haan, Maciej Grajewski, Patty P. M. F. A. Mulder, Jean-Paul S. H. Mulder, Hiddo J. Lambers-Heerspink, Elisabeth M. J. Verpoorte

## Abstract

Organ-on-a-chip (OOC) technology is a valuable tool that can facilitate a faithful replication of target tissues or organs by enabling the application of important physiological cues that cells experience in their native environment. We describe here an endothelium-on-a-chip that was developed for studying endothelial barrier permeability in diabetic and diabetic kidney disease (DKD) milieu under physiologically relevant shear stress. The fluorescently labelled macromolecular tracer flux assay was applied to obtain apparent permeability (P_app_) values. We demonstrated the protective effect of shear stress in our EoaC in diabetic and DKD milieu by testing endogenous diabetic and DKD factors individually (hyperglycaemia, pro inflammatory factor TNFɑ, and hypertension-related hormone aldosterone) and in combination (aldosterone in hyperglycaemia, as well as a DKD mix consisting of TNFɑ and aldosterone in hyperglycaemia). To assess the applicability of our microfluidic EoaC for drug testing, we added the mineralocorticoid receptor antagonist, finerenone, and observed a statistically significant protective effect of this pharmaceutical on endothelial barrier integrity in DKD milieu in our model system. We conclude that this EoaC can be applied for endothelial barrier permeability evaluation not only in diabetic and DKD milieu, but in various pathophysiological situations. Moreover, it has shown a potential to be used in drug testing targeted to improve endothelial barrier resilience.

## Introduction

Increasing demand for alternatives to animal testing has driven the development of *in vitro* test systems that are capable of recapitulating the physiology and pathophysiology of different organs and organ systems. Several examples of endothelium-on-a-chip (EoaC) exist for studying various aspects of the endothelial cell biology, for instance, cell adhesion and migration, angiogenesis, shear stress response, interactions with other cell types, as well as one of the main functions of endothelium, permeability.^1–5^ EoaC devices intended for permeability studies were initially designed by incorporating a porous membrane as the endothelial cell growth surface (as in the transwell approach) and measuring the passage of tracer molecules from the endothelial apical to basolateral side.^6–8^ Another approach involves the incorporation of the electrodes in the chambers on both sides of the membrane for transendothelial resistance measurements (TEER) (reviewed in ^9^). Both of these approaches have allowed researchers to evaluate changes in endothelial permeability under desired test conditions with application of shear stress in a more *in vivo*-like environment, compared to static systems. Further advancements striving to mimic the *in vivo* environment even better in an EoaC were developed in the last decade by incorporating ECM protein hydrogels as a growth surface for endothelial cells.^10–14^ The ECM protein mainly used for this purpose is collagen. A fluorescently labelled tracer is applied then on the apical side of the cells, for instance, the albumin or dextrans, coupled to fluorescein isothiocyanate (FITC). The diffusion of tracer into the ECM hydrogel is then monitored and quantified.

Diabetic kidney disease (DKD) is a leading cause of chronic kidney disease (CKD) worldwide.^15^ DKD develops in up to 40% of type 2 diabetes patients.^16^ It has a complex pathogenesis which is induced by metabolic, hemodynamic (pertaining to blood flow and vascular resistance), inflammatory and fibrotic (related to tissue thickening and scarring) factors (reviewed in ^17^). From a metabolic perspective, the hyperglycaemia induces the production of advanced glycation end products (AGEs) and reactive oxygen species (ROS) in cells. Changes in the regulation of the renin-angiotensin-aldosterone system (RAAS) increase angiotensin II and aldosterone levels and lead to mineralocorticoid receptor (MR) overactivation. All these key mechanisms contribute to the production of inflammatory and fibrotic factors and drive the progression of DKD. This leads to altered kidney filtration and barrier function, tubular damage, and even end stage renal disease (ESRD). In the context of DKD, developing an EoaC for assessing albumin leakage and evaluating the effects of the pharmaceuticals would facilitate more thorough characterisation of the kidney permeability barrier. The endothelial layer is the very first line of defence in the kidney, and an important regulator of permeability. Surprisingly, endothelial permeability in an EoaC for DKD research has not received much attention yet. Understanding the endothelial barrier permeability changes in DKD could be applied in the development of more targeted therapies for DKD patients, ultimately fostering strategies to tailor medications for individual patients, based on individual drug responses.

The knowledge generated in the study of the molecular mechanisms of DKD has been applied to develop therapies for patients with DKD, including angiotensin converting enzyme inhibitors (ACEi), angiotensin receptor blockers (ARBs), sodium-glucose cotransporter-2 (SGLT2) inhibitors and the nonsteroidal mineralocorticoid receptor antagonist (MRA) finerenone.^18^ Finerenone has been shown to delay the progression of DKD and reduce the risk of cardiovascular events (reviewed in ^19^). Finerenone is a third-generation MRA, which is highly selective with strong affinity for the MR. In contrast to steroidal MRAs (e.g. spironolactone), finerenone exhibits a lower risk for hyperkalemia and androgen-like effects.^20,21^ Finerenone has been shown to reduce the urinary albumin-to-creatinine ratio (UACR) in patients with DKD.^22^ The early change in UACR is associated with subsequent cardio-renal risk,^23^ suggesting that the reduction in UACR may be used as a therapeutic efficacy marker of the finerenone. Another group of pharmaceuticals, endothelin receptor antagonists (atrasentan, sparsentan and others), have received ample attention in recent years. They have been applied for the treatment of CKD, based on the findings that these pharmaceuticals reduce albuminuria and the risk of kidney failure.^24,25^

We report here a development and characterisation of an EoaC for endothelial barrier layer permeability studies and modelling the milieu typical to DKD by application of endogenous DKD factors individually and in combination. Additionally, we evaluated the suitability of this EoaC for application in drug research. Our polydimethylsiloxane (PDMS) EoaC has a simpler design for permeability evaluation upon application of flow in comparison to previously reported EoaC devices. The ECM hydrogels in EoaC devices are usually introduced in a channel that lies parallel, on the same plane, to the endothelial cell growth channel.^12^ This means that the endothelial cells need to form a monolayer on a vertical “wall” of the ECM gel. To facilitate endothelial cell settling and adherence on the ECM hydrogel “wall”, these EoaC devices have to be tilted by 90° after cell seeding for a certain amount of time. In our EoaC, the hydrogel is located on an x-z plane, forming the floor of a flow channel. The endothelial cells settle and adhere to the hydrogel by gravity. This means that no additional device rotation step is necessary after cell seeding to establish a cell layer. This reduces disturbances on the endothelial layer in the EoaC and ensures optimal conditions for the endothelial cells to rapidly reach confluency.

We have chosen to work with gelatin as the endothelial cell growth surface, as it is a naturally occurring ECM that can be cross-linked with microbial transglutaminase (mTG), an enzyme that is non-toxic for cells.^26,27^ Gelatin is a derivative of collagen and is economically more convenient.^28^ We have utilised this hydrogel previously in our “proof-of-concept” experiments in well plates in static conditions in diabetic and DKD milieu, and shown that it allows quantification of a permeability tracer that has crossed the endothelial barrier (unpublished; manuscript in preparation).

We characterised the EoaC in terms of endothelial cell morphology, as well as functionality regarding permeability. We then modelled the diabetic and DKD environment in an EoaC upon application of physiologically relevant shear stress (10 dyn/cm^2^). Finally, we tested the pharmaceutical finerenone, targeted to endothelial permeability resilience, to evaluate the potential of developed EoaC to be applied in drug testing.

### Experimental

#### General layout of the EoaC device

The EoaC device has two PDMS layers and a glass microscope slide on the bottom (Fig. 1). The top PDMS layer has a microfluidic channel that supplies endothelial cell medium and test factors to the cells. The lower PDMS layer hosts a channel for the gelatin/mTG hydrogel.

**Fig. 1.**
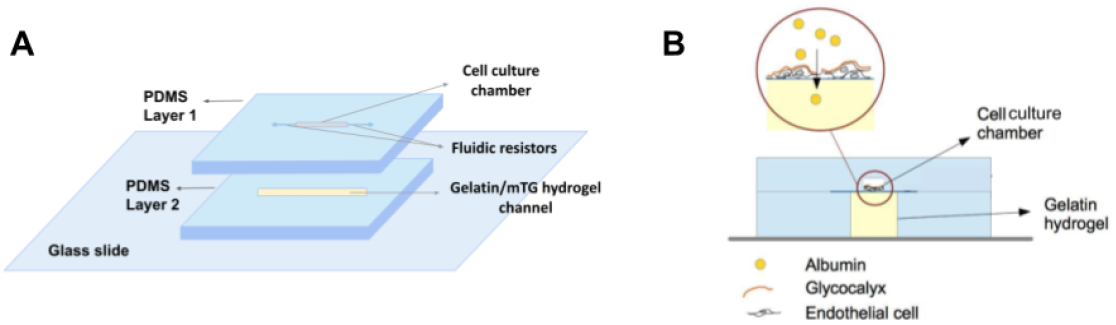
Layout and cross-sectional view of the EoaC device. A - the EoaC device consists of two PDMS layers and a glass slide on the bottom. Layer 1 has a cell culture chamber with fluidic resistors. It is used for cell seeding and for delivery of endothelial cell medium and test factors. Layer 2 has a cavity for gelatin/mTG hydrogel; B - a cross-sectional view of the EoaC explaining the working principle of the fluorescence-based endothelial permeability assay. Gelatin/mTG hydrogel has two functions in our EoaC. First, it acts as a biomimetic growth surface for endothelial cells. Second, it receives the permeability tracer, albumin-FITC, which passes the endothelial barrier layer and collects in the hydrogel. The analysis of albumin-FITC content in the gelatin/mTG hydrogel allows to characterise the permeability properties of the endothelial barrier.

The top PDMS layer has a flow microchannel with a 10-mm-long, 0.36-mm-wide and 0.1-mm-high cell cultivation chamber and two 5-mm-long, 0.18-mm-wide and 0.1-mm-high fluidic resistors (Fig. 1 A). The shape of the flow microchannel was selected according to the findings of Grajewski et al.^29^ It has been observed that this channel geometry ensures longer retention of the cells in the cell cultivation chamber after cell seeding, as the inlet resistor slows down the flow of the cell suspension, and the outlet fluidic resistor adds a backpressure. This results in higher cell numbers and, consequently, a higher initial cell confluency in the EoaC. High confluency of the cell layer in the EoaC device is crucially important for barrier permeability studies. For this reason, we strive for cell confluency as close as possible to 100% in the EoaC.

We incorporated a gelatin/mTG hydrogel in the lower PDMS layer of the EoaC device to act as an ECM. The presence of ECM in cell cultures has been shown to ensure more relevant cell-cell and cell-ECM interactions.^30^ The application of flow in the EoaC adds another crucial physiological factor for endothelial cells. It imitates the blood flow and recreates the effects of shear stress experienced by the endothelial cells *in vivo*. The gelatin/mTG hydrogel in our EoaC has an additional role in the fluorescence-based permeability assay. It acts as a receiving phase for the fluorescently labelled permeability tracer, albumin-FITC, after it has passed endothelial barrier from the apical side and diffused into the hydrogel (Fig. 1 B).

After exposure to the test factors in the EoaC, the endothelial cell layer is incubated with albumin-FITC. Normally, the majority of albumin is prevented from crossing the endothelial barrier due to its strictly regulated permeability.^31^ A decrease of endothelial barrier integrity due to the damaging effects of test substances will result in an increased permeability. Consequently, increased concentrations of albumin-FITC will be able to pass the endothelial barrier and diffuse into the gelatin/mTG hydrogel. This will manifest as a higher fluorescence intensity of albumin-FITC in the gelatin/mTG hydrogel. We then correlate the fluorescence intensity measurements of the albumin-FITC in the hydrogel with transferred albumin-FITC concentration and calculate apparent permeability (P_app_) values.

#### Casting a gelatin/mTG hydrogel channel in PDMS

To prepare the gelatin/mTG hydrogel channel in PDMS, a micro-milled polymethylmethacrylate (PMMA) positive mould was used (Fig. S 1, Supporting Information). PDMS elastomer (Sylgard 184, Dow Corning Corp., MI, USA) was mixed with a curing agent (Sylgard 184, Dow Corning Corp., MI, USA) at a 10:1 ratio, degassed for 30 min at room temperature (RT) and poured in the PMMA mould. The PMMA mould with PDMS was then cured on a hotplate (Harry Gestigkeit GmbH, Germany) at 70°C for 1.5 h. After, it was manually peeled from the PMMA mould. The resulting PDMS layer had a 25-mm-long, 3-mm-deep and about 0.5-mm-wide channel for the gelatin/mTG hydrogel.

PDMS is an elastic material, which can be a challenge in the fabrication procedure. The lower PDMS layer that contains the hydrogel can be deformed accidentally, thereby leading to damage to the hydrogel in the channel (undesired tears or artefacts). To mechanically support the lower PDMS layer before introducing the hydrogel, the bottom surface of it was treated in an oxygen plasma cleaner (Harrick Plasma Cleaner; Harrick Plasma, NY, USA) at ∼ 360 mTorr for 20 s. The glass microscope slide (Gerhard Menzel & Co., Germany) was treated the same way, and both parts were irreversibly bonded by firmly pressing together the treated sides of the PDMS and the glass slide (Fig. S 1 A, Supporting Information).

A PDMS cover sheet was prepared to contain the gelatin/mTG hydrogel and to ensure a flat and even surface of the gelatin hydrogel for cell growth. It was prepared by pouring 20 g of PDMS elastomer-curing agent solution (10:1) in a rectangular Petri dish (120 x 120 mm) (VWR, PA, USA) and curing it at 70°C for 1.5 h. After curing, the PDMS cover sheets were cut with a scalpel to the desired size (length ∼ 4 cm, width ∼ 1.1 cm), and inlet and outlet holes were punched with a 2-mm biopsy puncher (Kai Medical, Japan) (Fig. S 1 B, Supporting Information). The approximate thickness of the PDMS cover sheets was 0.15 cm. The inlet and outlet were positioned to align with the ends of the gelatin hydrogel channel. The PDMS cover sheet was then placed on top of the PDMS hydrogel channel such that the inlet and outlet matched with the gelatin/mTG channel (Fig. S 1 C, Supporting Information). The lower PDMS layer was now ready for injecting gelatin/mTG solution.

Storage of it is possible at RT in such a way that prevents the dust from entering the hydrogel channel, for instance, by placing it in a covered Petri dish.

#### Gelatin/mTG hydrogel preparation in the hydrogel channel

Gelatin/mTG hydrogel was prepared following the protocol of Paguirigan and Beebe with slight modifications.^28^ Briefly, a solution of gelatin with a final concentration of 10% was prepared; 10 U microbial transglutaminase (mTG) (activity > 30 U/g, ASA Spezialenzyme GmbH, Germany) per gram of gelatin was added to this solution. For 10 mL of hydrogel, gelatin solution and mTG solution were prepared separately. Gelatin solution was prepared by dissolving 1 g gelatin (Type A from porcine skin, ∼300 Bloom, Sigma-Aldrich, Germany) in 8 mL of sterile 1x PBS (pH 7.4, Gibco, Thermo Fisher Scientific, OR, USA). Gelatin solution was heated at 50°C on a hotplate for 30 min to dissolve gelatin. This was followed by additional heating at 55°C for 15 min, occasionally swirling the solution. Enzyme solution was prepared by diluting 0.33 g mTG in 0.67 mL sterile 1x PBS. Enzyme solution was then added to the gelatin solution and mixed thoroughly. Immediately after mixing, gelatin/mTG solution was injected into the hydrogel channel through the inlet of the PDMS cover sheet using a 1-mL syringe without needle (BBraun, Germany). The excess gelatin/mTG solution escaped through the outlet in the PDMS cover sheet. It is essential to avoid air bubbles and work fast, as the gelatin/mTG solution becomes very viscous in about two minutes.

The gelatin/mTG hydrogel in the channel was crosslinked at 37°C for 3 h. Afterwards, the PDMS layer with gelatin/mTG hydrogel was transferred to the hotplate and incubated at 65°C for 30 min to heat-inactivate the enzyme. The PDMS cover sheet was carefully removed with tweezers and replaced with another temporary PDMS sheet to protect the gelatin/mTG hydrogel from drying during the oxygen plasma treatment. The temporary PDMS sheet was prepared by pouring 20 g of PDMS elastomer-curing agent solution (10:1) in a rectangular Petri dish and incubating it at 70°C for 1.5 h. The resulting temporary PDMS sheet was cut out with a scalpel to a desired size (length ∼ 3 cm, width ∼ 0.5 cm) to cover the hydrogel-filled channel completely. The thickness of the temporary PDMS cover sheet was approximately 0.15 cm. The temporary PDMS sheet fully covered the gelatin/mTG hydrogel and slightly overlapped with the edges of the PDMS layer. We selected the size of the temporary PDMS cover sheet to obtain sufficient hydrogel protection during the oxygen plasma treatment and to leave as much as possible of the PDMS surface exposed to oxygen plasma. This ensured strong bonding with no leaks between the PDMS layers.

#### Preparing the flow microchannels in PDMS

Designing of the microchannel, the preparation of SU-8 50 master mould and casting the microchannels in PDMS was performed according to the method described in the Supporting Information.

#### Assembly of the EoaC device and preparation for cell seeding

Both PDMS layers of the EoaC device were treated in the oxygen plasma cleaner at ∼ 360 mTorr for 20 s to create the irreversible bond. The hydrogel channel and the flow channel were aligned with an aligning tool (Edmund Optics, Japan), and brought into contact with a gentle pressure to form an irreversible bond. To ensure an aseptic environment, all further manipulations with the EoaC devices were performed in the laminar air hood (Nino Labinteriör AB, Sweden).

To sterilise the surface of the hydrogel, 70% ethanol was introduced into the microchannels with a 1-mL syringe without needle, and incubated at RT for 30 min. The microchannels of EoaC devices were then rinsed three times with sterile 1x PBS by introducing the 1x PBS through the inlet and aspirating it through the outlet with a 1-mL syringe without needle. The EoaC devices were then preconditioned by introducing endothelial cell culture medium (Endothelial Growth Medium-2 (EGM-2) composed of Endothelial Basal Medium-2 (EBM-2) with added SingleQuots^TM^ Supplements (Lonza, Switzerland) in the microchannels and incubating overnight at 37°C. This allowed for the saturation of gelatin/mTG with nutrients from the endothelial culture medium.

Cell maintenance and seeding in the EoaC device is described in the Supporting Information.

#### Flow system setup: components and application of the flow to the EoaC

The flow setup for EoaC consists of several components: syringe pumps; syringes; connectors for syringes and inlet tubes; inlet and outlet tubes; and waste containers (for a detailed description, see Supporting Information and Fig. S 2).

Sterile 60-mL syringes were filled with endothelial cell culture medium and incubated at 37°C for about two hours prior to addition of flow. This step reduced air bubbles in the cell culture medium and pre-warmed it before coming into contact with the cells. Inlet tubes were connected to the syringe as described, and filled with endothelial cell medium from the syringe. The inlet and outlet tubes were inserted in the EoaC inlet and outlet, respectively. The other end of the outlet tube was inserted in a sterile waste container of appropriate volume.

We used laminar unidirectional flow in our experimental setup. A physiologically relevant shear stress of 10 dyn/cm^2^ was applied to our EoaC. The shear stress was selected based on literature data for relevant shear stress for endothelial cells in the kidney, which ranges from 5 to 20 dyn/cm^2^.^32^ To obtain 10 dyn/cm^2^ shear stress, we calculated the necessary flow rate using Equation 1:^33^

**Equation 1** Equation for calculating flow shear stress.

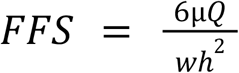

where **FFS** - flow shear stress (dyn/cm^2^);

**µ -** viscosity of endothelial cell medium (ca. 0.0075 dyn*s/cm^2^);^29^

**Q** - flow rate (cm^3^/s);

**w** - width of the microfluidic channel (cm);

**h** - height of the microfluidic channel (cm).

According to the equation, we set the flow rate to 46.11 µL/min to obtain shear stress of 10 dyn/cm^2^ in our EoaC. After the experiment, the inlet and outlet tubes were removed and fluorescence-based permeability assay was performed in the EoaC.

#### Modelling DKD milieu in an EoaC for permeability assay and evaluation of pharmaceuticals

Several endogenous DKD factors separately or in combinations were selected for testing in endothelium-on-chips (EoCs) with application of 10 dyn/cm^2^ shear stress. The concentrations of these endogenous diabetes and DKD factors were selected in accordance with the literature data. For testing in EoaC, we selected the elevated glucose (33 mmol/L),^34–36^ proinflammatory cytokine TNFɑ (10 ng/mL)^37–39^ and hormone aldosterone (10 nmol/L)^40^ individually. The cell culturing media contains 5.55 mmol/L glucose, therefore representing a physiologically normal glucose concentration. Since diabetic and DKD conditions are complex and have several endogenous factors present simultaneously, we also tested the combination of aldosterone (0.1 nmol/L)^41^ in hyperglycaemia (33 mmol/L), and a DKD mix (glucose: 33 mmol/L; aldosterone: 0.1 nmol/L; and TNFɑ 10 ng/mL). We added the test factors to the endothelial cell medium. Three biological replicates (three EoCs) were prepared for each test condition.

We stimulated cells for 6 h (hyperglycaemia and aldosterone individually) or for 24 h (TNFɑ individually; the combination of aldosterone in hyperglycaemia) under shear stress of 10 dyn/cm^2^ at 37°C in 5% CO_2_. When testing DKD mix, we first incubated an EoaC with shear stress of 1 dyn/cm^2^ for about 72 h at 37°C in 5% CO_2._ We then stimulated the cells in the EoaC with DKD mix or DKD mix with added finerenone (1µM) under 10 dyn/cm^2^ shear stress for 24 h at 37°C in 5% CO_2_. The working concentration of finerenone was determined according to the method described in Supporting Information.

After endothelial cell stimulation, the flow system tubes were removed and a permeability assay was carried out in the EoaC under no-flow conditions.

#### Fluorescence-based permeability assay in an EoaC

Albumin-FITC (3 µM) (Sigma-Aldrich, Germany) was introduced through the inlet in the flow microchannel of EoaC using a micropipette with a 100-µL pipette tip (Gilson, WI, USA). Another 100-µL pipette tip was inserted in the outlet of EoaC to collect the excess albumin-FITC solution. This procedure was carried out in a darkened laminar flow hood to protect the samples from the background light that might initiate photobleaching. The EoaC with inserted inlet and outlet pipette tips was then incubated in the dark at 37°C for 30 min without flow. The inserted pipette tips helped to contain the fluorescent albumin-FITC. After incubation, 100-µL pipette tips were removed from inlet and outlet. The microchannel was rinsed with micropipette and 100-µL micropipette tips three times with 100-µL sterile 1x PBS. The excess PBS was collected in a 100-µL pipette tip in the outlet, which was exchanged after each rinsing. After rinsing, a cell culture medium was introduced in the EoaC and the imaging was performed. Fluorescence and light microscopy images were obtained.

One-point calibration was carried out to correlate the measured fluorescence intensity of albumin-FITC in the hydrogel to the known concentration of albumin-FITC (3 µM). The EoaC device for one-point calibration without cells was prepared for each experiment. The one-point calibration devices were prepared along with the experimental EoaC devices and experienced the same timeline as experiment devices. The only exception was that flow was not applied to the one-point calibration microfluidic devices. When performing the permeability assay, the one-point calibration images were taken immediately after introducing 3 µM albumin-FITC into the channels of the device. One-point calibration was selected to enable more quantitative determination of the concentrations of albumin-FITC that has passed the endothelial barrier layer and diffused in the gelatin/mTG hydrogel underneath. Calibration compensates for slight variations in fluorescence intensity that can manifest from one experiment to the next due to changes in background light or other environmental factors (e.g., temperature, etc.).

#### Imaging and result analysis

The EoaC was imaged with an inverted microscope Leica DMIL LED (light source: EL6000; camera: Leica DFC310 FX; objective magnification: 5x; and aperture: 0.12) (Leica, Germany). The method of taking the images is described in detail in the Supporting Information.

The mean value of the corrected fluorescence intensity measurements in a.u. for each EoaC was calculated from seven measurements. Three biological replicates were obtained. As a result, 21 fluorescence intensity measurements were obtained for each test condition. The conversion factor (fluorescence intensity : albumin-FITC concentration ratio) was calculated from a one-point calibration. This allowed us to calculate the albumin-FITC concentration that had diffused into the gelatin/mTG hydrogel (Q).

The apparent permeability, P_app_, was calculated with Equation 2 (from ^12^ and ^42^):

**Equation 2** Equation for calculating P_app_.

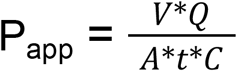

where **P**_**app**_ is apparent permeability (cm/s);

**Q** is the concentration of albumin-FITC diffused in the hydrogel (mM);

**V** is the volume of the hydrogel (0.0375 cm^3^);

**A** is the surface area of the hydrogel (0.125 cm^2^);

**t** is the time (1800 s);

**C** is the initial concentration of albumin-FITC (mM).

Box and whiskers graphs were then prepared with the program Prism 9 (Version 9.2.0, GraphPad Software, CA, USA). The statistical significance of the results was determined using analysis of variance (ANOVA) test followed by a post-hoc comparison of each test condition to the respective control (ANOVA with Dunnett’s test).

## Results and discussion

### Characterisation of the EoaC device

We designed and manufactured a simple two-layer PDMS EoaC incorporating ECM hydrogel of natural origin, gelatin/mTG hydrogel. We prepared the PDMS layers of our EoaC device with a soft lithography technique. The resulting device was bonded to a glass microscope slide and was both transparent and gas permeable. The volume of the EoaC microchannel was calculated to be 0.54 µL. The transparency of the EoaC device allowed cell monitoring by microscopy. Light or fluorescence microscopy techniques could be applied to monitor the endothelial layer in the EoaC. In our observations, PDMS and the gelatin/mTG hydrogel exhibited practically no autofluorescence. The design of this EoaC allows easy application of a range of shear stresses simply by varying the cell medium flow rate. The EoaC device was leakage-proof and biocompatible as confirmed by observation of endothelial cell settlement on gelatin/mTG hydrogel, followed by attachment and monolayer formation.

### Endothelial cell morphology in an EoaC

Permeability studies require a confluent endothelial cell layer to begin with. We observed endothelial monolayer formation as soon as one hour after HUVEC seeding in the EoaC (Fig. 5 A). The cell confluency three hours after cell seeding was close to 100%, as observed by cell microscopy.

**Fig. 5.**
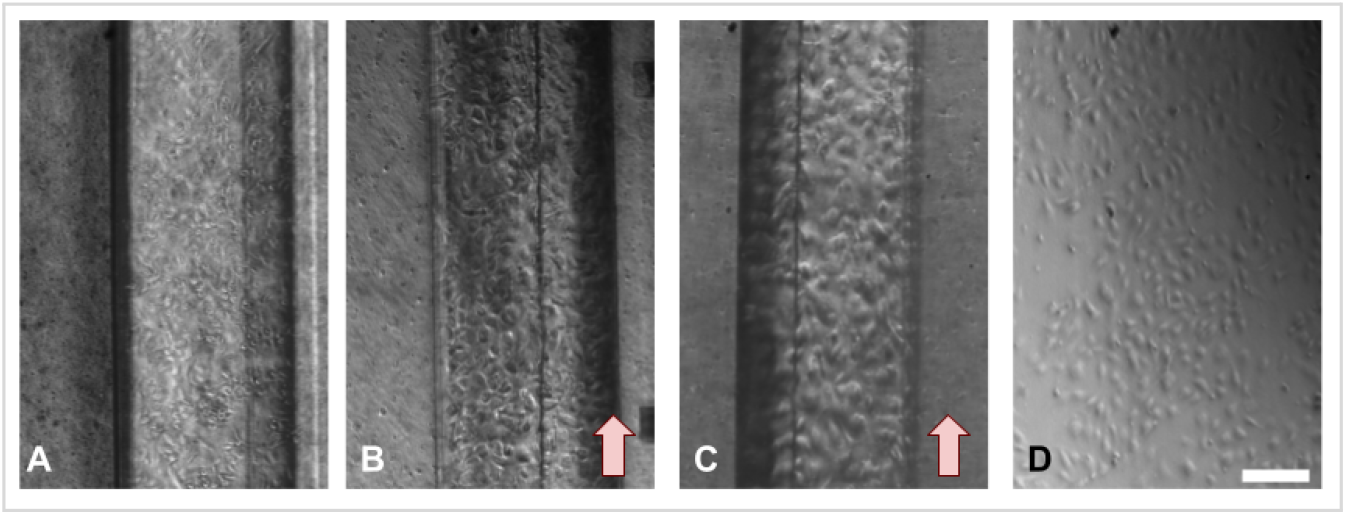
Endothelial cell morphology in an EoaC. A - HUVECs one hour after seeding in an EoaC. Monolayer formation can be observed; B - HUVECs after 24 h of exposure to shear stress (10 dyn/cm^2^) in an EoaC. Cells have a typical cobblestone-like morphology; C - HUVECs after approx. 96 h of exposure to shear stress (1 dyn/cm^2^ for the first 72 h and 10 dyn/cm^2^ for the following 24 h). HUVECs maintained their typical cobblestone-like morphology; D - HUVECs cultured for 24 h in a cell culturing flask in static conditions. Typical cobblestone-like cell morphology can be observed. The arrows indicate flow direction. Scale bar: 200 µm.

After 24 h of HUVEC culturing in the EoaC with applied shear stress (10 dyn/cm^2^), we observed a typical cobblestone-like cell morphology (Fig. 5 B), similar to that of static control (Fig. 5 D). Endothelial cells maintained their typical morphology after approximately 96 h of culturing with applied shear stress (1 dyn/cm^2^ for the first ∼ 72 h and 10 dyn/cm^2^ for the following 24 h) in our EoaC (Fig. 5 C).

Interestingly, our observations about endothelial cell morphology in response to shear stress do not align well with the literature data. There is a large amount of evidence in the literature pointing to HUVEC elongation in the direction of flow under similar shear stress conditions. For instance, Bertani and colleagues observed HUVEC alignment and elongation after 24 h under 10 dyn/cm^2^ shear stress along with elevated actin stress fibre expression at the cell periphery.^43^ Similar observations were made by Grajewski and colleagues after just 20 h of HUVEC culture under 10 dyn/cm^2^ shear stress.^29^ Reinitz *et al*. observed HUVEC elongation after 36 h in 12 dyn/cm^2.44^ Another research group, Silvani *et al*., noticed HUVEC alignment in the flow direction at a shear stress of 16 dyn/cm^2^ after 30 h.^45^ The discrepancy between our results and the results in the literature might be attributed to the differences in the stiffness of the cell growth surface. The studies mentioned above utilised cell growth surfaces coated with fibronectin ^44–46^ or glutaraldehyde-crosslinked gelatin.^29^ ECM coatings have different tensile properties (Young’s modulus) compared to the gelatin/mTG hydrogel. For instance, the Young’s modulus for the fibronectin coating has been reported generally to be in the range of MPa, whereas this parameter falls in the range of kPa for gelatin/mTG hydrogel.^47^ This means that the gelatin/mTG hydrogel used in our EoaC provides a far less stiff substrate for endothelial cell growth compared to surface coatings.

Changes in cell morphology, contractility and cell-cell and cell-ECM interactions as a response to the growth surface stiffness have been well documented by a number of studies on various cell types, including endothelial cells (reviewed in ^48^). Generally, cell morphology is dictated by the dynamics of the cell cytoskeleton, which responds to a range of physiological and chemical cues. It has been hypothesised by Ingber *et al*. that the cytoskeleton is stabilised by a force balance between the tensile stress experienced by actin microfilaments and compression forces of the intracellular environment supporting structures (microtubules and intermediate filaments) as a result of interactions from outside of the cell (traction forces due to binding to the ECM through focal complexes, and to the neighbouring cells).^49^ This concept is called tensional integrity (or tensegrity). Changes in HUVEC morphology under shear stress have been explained in the literature as being due to a reorganisation of actin stress filaments along the cell periphery and remodelling of intercellular junctions, as well as intermediate filament reorganisation.^44–46,50^ In our case, we hypothesise that the intracellular interactions (between actin microfilaments, microtubules and intermediate filaments) and intercellular (cell-cell) interactions of the endothelial cells were able to compensate for the extracellular traction forces applied on cells. These extracellular forces are due to the stiffness of the gelatin/mTG hydrogel in combination with shear stress (Fig. 6).

**Fig. 6.**
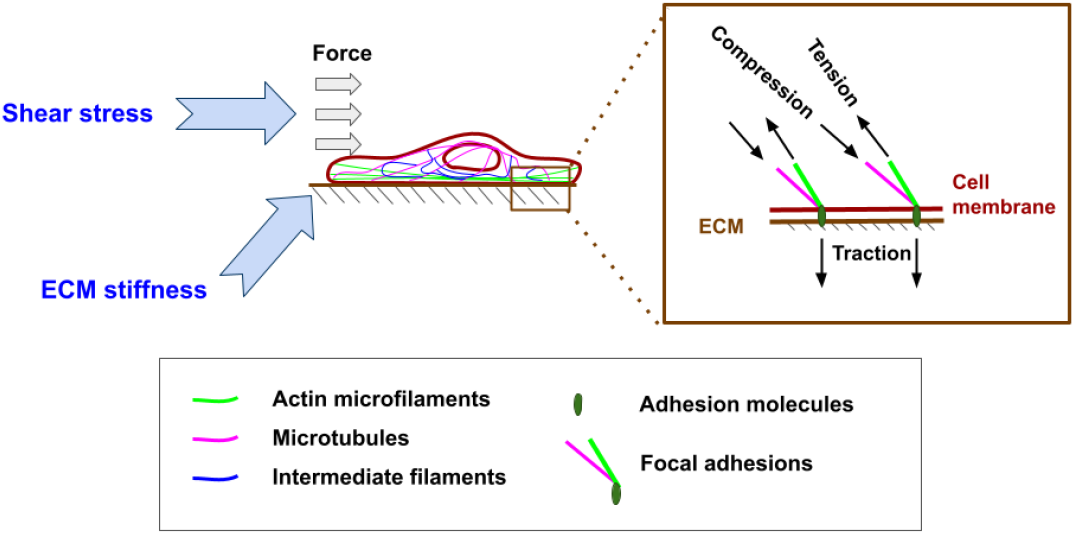
The external forces applied on endothelial cells in our EoaC. We hypothesise that the balance between the effects of external forces (shear stress and stiffness of the gelatin/mTG hydrogel) on actin microfilaments and opposing forces of intracellular support structures (microtubules and intermediate filaments), as well as cell-cell interactions, maintain the typical cobblestone-like endothelial cell morphology under shear stress (a concept of tensional integrity).^49^

Moreover, the Young’s modulus of the gelatin/mTG hydrogel has been reported to be 93.0 ± 24.1 kPa, which is close to the Young’s modulus of human kidney. The Young’s modulus of human kidney has been shown to range from 5-10 kPa for the whole kidney to 95 kPa (transversal loading) or 180 kPa (axial loading) for cut kidney tissue.^51–53^ The fact that our gelatin/mTG hydrogel has similar tensile properties as human kidney tissue ensures more relevant cell-ECM interactions. This allows us to model the DKD environment in our EoaC with a closer resemblance to the *in vivo* environment.

### Endothelial layer permeability in the EoaC

We applied the fluorescently labelled tracer, albumin-FITC, to characterise permeability properties of the endothelial barrier layer in the EoaC. Our EoaC facilitates the quantification of albumin-FITC that has passed through the endothelial barrier and diffused into the gelatin/mTG hydrogel. The calculated mean P_app_ for HUVECs cultured in the EoaC for 24 h under a shear stress of 10 dyn/cm^2^ was 8.32 × 10^-5^ ± 1.89 × 10^-5^ cm/s (for three biological replicates). This value is about one order of magnitude higher than what we have obtained for HUVECs in similar conditions in a static culture in our earlier experiments (8.25 × 10^-6^ ± 0.93 × 10^-6^ cm/s, from one biological replicate) (unpublished; manuscript in preparation). We thus observe an increase in the endothelial layer permeability after application of shear stress to the endothelial cells. A temporary increase in endothelial layer permeability due to application of the shear stress has been observed by many previous *in vitro* studies. For instance, Warboys *et al*. observed an increased endothelial permeability after a 1-h exposure to the shear stress; however, exposure to shear stress for seven days decreased the endothelial permeability, compared to a static control.^54^ The authors indicated that this result demonstrates the effects of acute and chronic shear stress on endothelial cell permeability and that this indicates the resilience of the endothelial barrier. Another research team studied the effects of 10 dyn/cm^2^ shear stress on endothelial cells after only 1 h, and, as well, observed dramatically increased endothelial permeability.^55^ The permeability returned to pre-shear-stress values after removal of the shear stress. Taken together, these studies, along with our results, highlight the dynamic nature of the endothelial barrier layer in the presence of shear stress.

### The effects of DKD environment on endothelial barrier layer in an EoaC with application of shear stress

#### Hyperglycaemia and TNFɑ triggers changes in endothelial cell morphology in an EoaC with applied shear stress

We observed HUVEC morphology under shear stress after challenging the cell monolayer for 6 or 24 h in our EoaC with the factors present in diabetes and DKD. The endothelial cell morphology in the control EoaC exhibited typical cobblestone-like morphology after 6 and 24 h of application of 10 dyn/cm^2^ shear stress (Fig. 7 A and D).

**Fig. 7.**
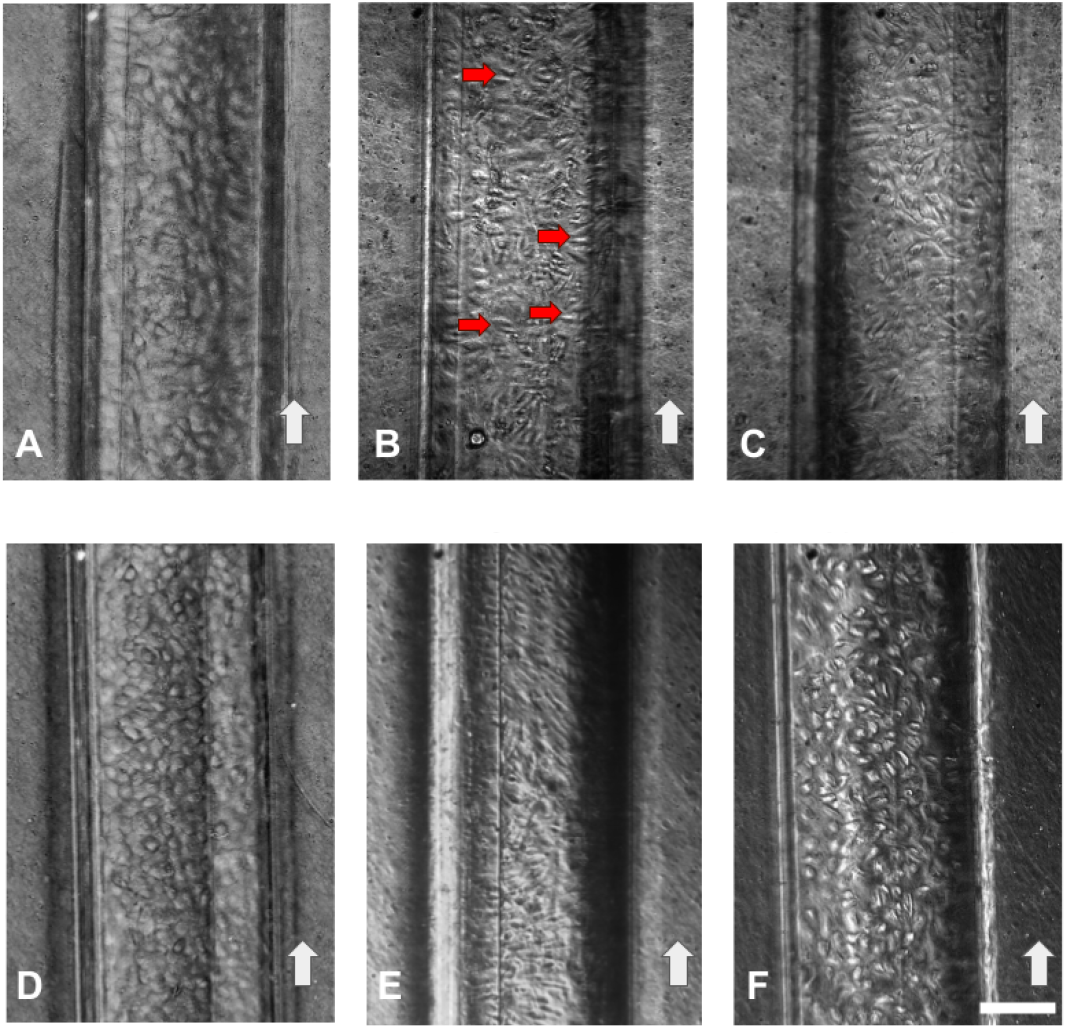
Endothelial cell morphology in an EoaC with an applied shear stress of 10 dyn/cm^2^ after exposure to DKD factors. A - control HUVECs after 6 h of application of shear stress; B - HUVECs after 6 h of stimulation with elevated glucose (33 mmol/L) and application of shear stress. Red arrows indicate areas with changed (elongated) cell morphology; C - HUVECs after 6 h of stimulation with aldosterone (10 nmol/L) and application of shear stress; D - control HUVECs after 24 h of application of shear stress; E - HUVECs after 24 h of stimulation with TNFɑ (10 ng/mL) and application of shear stress; F - HUVECs after 24 h of stimulation with aldosterone (0.1 nmol/L) in hyperglycaemia (33 mmol/L) and application of shear stress. The white arrows indicate the direction of the flow. Scale bar: 200 µm.

Slight changes in endothelial morphology could be observed after cell exposure to hyperglycaemic conditions (33 mmol/L) for 6 h (Fig. 7 B). Some of the HUVECs seem to have become slightly elongated (indicated with red arrows). It is well known that the hyperglycaemic environment triggers the formation of the reactive oxygen species (ROS) and thus oxidative stress in endothelial cells.^56^ Oxidative stress in endothelial cells have been previously associated with cell elongation and actin stress fibre formation, which aligns with our observations.^57^

Pronounced endothelial cell elongation was observed in the EoaC after cell stimulation with proinflammatory cytokine TNFɑ (10 ng/mL) for 24 h with applied shear stress (10 dyn/cm^2^) (Fig. 7 E). This result is in good agreement with literature data. TNFɑ is a potent inflammatory trigger, and a large volume of evidence in the literature demonstrates its effect on endothelial cell morphology. For instance, Stroka and colleagues observed strongly elongated HUVECs after 8 h and 24 h of stimulation with 25 ng/mL TNFɑ in a static system along with increased traction forces generated by HUVECs, compared to control cells.^58^ Wang and colleagues demonstrated concentration-dependent effects of TNFɑ on HUVEC morphology.^59^ These morphological changes have been associated to a TNFɑ-induced actin reorganisation and stress fibre formation through Rho and myosin light chain kinase (MLCK) dependent mechanisms.^60^ The same mechanisms have been found to be partly involved in permeability increase in endothelial cells after stimulation with TNFɑ.^61,62^

#### Shear stress has a protective effect on endothelial layer integrity in DKD milieu

To evaluate the response of the endothelial cell layer to endogenous DKD stimuli under shear stress in our EoaC, we stimulated HUVECs for 6 or 24 h with DKD factors individually or in combination.

After 6 h of endothelial cell incubation with glucose (33 mmol/L) and the hormone aldosterone (10 nmol/L) with application of 10 dyn/cm^2^ shear stress, we observed a decreased endothelial layer permeability, compared to the control (where glucose = 5.5 mmol/L). The mean P_app_ for the HUVEC layer stimulated with glucose (33 mmol/L) was 7.61 × 10^-5^ ± 0.99 × 10^-5^ cm/s, and for cells stimulated with aldosterone (10 nmol/L) 8.04 × 10^-5^ ± 0.42 × 10^-5^ cm/s, in comparison to mean P_app_ of the control of 8.99 × 10^-5^ ± 0.61 × 10^-5^ cm/s. These differences between DKD factors and control were found to be statistically significant (Fig. 8).

**Fig. 8.**
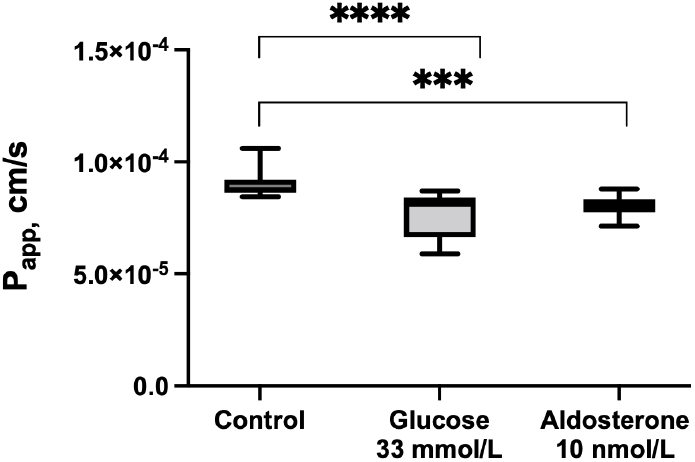
Endothelial permeability (P_app_) after 6 h of exposure to DKD factors with applied shear stress (10 dyn/cm^2^) in an EoaC. The result for each test condition was obtained from three biological replicates (n=3). Seven images were taken per biological replicate, and 21 images in total were analysed for each condition. Each box plot depicts a minimum and maximum value, as well as median and lower and upper quartiles of the data set. Level of significance indicated: *** - 0.0001 to 0.001; **** - < 0.0001.

We investigated the effects on HUVEC permeability in the EoaC after 24 h of stimulation with proinflammatory cytokine TNFɑ (10 ng/mL), and a combination of aldosterone (0.1 nmol/L) in hyperglycaemia (33 mmol/L) with application of shear stress (10 dyn/cm^2^) (Fig. 9). A slightly elevated endothelial permeability was observed on the barrier layer stimulated with TNFɑ (mean P_app_ 9.74 × 10^-5^ ± 1.53 × 10^-5^ cm/s) in comparison to the control (mean P_app_ 8.43 × 10^-5^ ± 1.63 × 10^-5^ cm/s); however, this effect was not statistically significant. In our earlier experiments in a static system, we observed a statistically significant (p < 0.0001) increase in endothelial layer permeability (unpublished; manuscript in preparation).

**Fig. 9.**
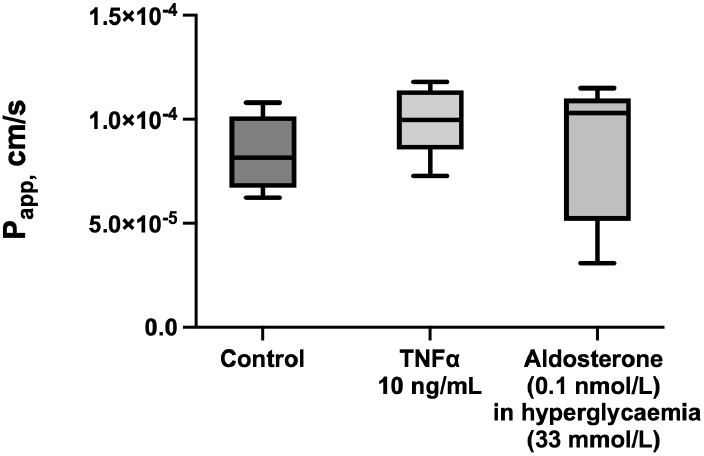
Endothelial permeability (P_app_) after 24 h of exposure to DKD factors with applied shear stress (10 dyn/cm^2^) in an EoaC. The result for each test condition was obtained from three biological replicates (n=3). Seven images were taken per biological replicate, and 21 images in total were analysed for each condition. Each box plot depicts a minimum and maximum value, as well as median and lower and upper quartiles of the data set.

The results from our experiments indicate a trend of a reduced effect of endogenous DKD factors on endothelial permeability after application of flow in comparison to a static system. It has been demonstrated previously that laminar flow within the physiological range is able to suppress inflammatory signalling in order to stabilise the vessels. Specifically, it has been shown that in HUVECs, the protective pathways are activated with the application of shear stress between 10 and 20 dyn/cm^2^.^63^ For instance, a dramatic increase in Krüppel-like factor 2 (KLF2) mRNA expression has been demonstrated after application of 10 dyn/cm^2^ shear stress for 24 h to conditionally immortalised human glomerular endothelial cells (CiGEnCs), in comparison to static cultures.^41^ KLF2 is a shear-stress-induced transcription factor that is involved in endothelial permeability regulation, and its overexpression tightens endothelial permeability barrier.^64,65^ In another example, a normalisation of phosphorylated endothelial nitric oxide (p-eNOS) levels in hyperglycaemia (33 mM) in comparison to normoglycaemia was shown after application of shear stress (20 dyn/cm^2^) for 24 h in a porcine endothelial cell culture.^66^ In this study, the phosphorylation of eNOS was associated with increased NO levels, which is a known contributor to increased vascular permeability.^67^

We hypothesise from this that the application of a shear stress of 10 dyn/cm^2^ in our EoaC has activated protective pathways in HUVECs and resulted in enhanced barrier resilience in DKD milieu. To explore this hypothesis, a deeper investigation should be conducted to find out whether these effects are transitional or chronic, and which pathways might be involved.

### Evaluating pharmaceuticals in a DKD EoaC

#### Finerenone does not have an impact on DKD-induced cell morphology

We evaluated the action of 1 µM finerenone in the DKD milieu on endothelial morphology under shear stress in our EoaC. We prepared three EoCs for each test condition, representing three biological replicates. However, due to the malfunction of one EoaC for the control and one EoaC for the DKD environment, we were only able to analyse two biological replicates (n=2) for each of these two test conditions. Ideally, three biological replicates should be considered for each test condition to increase the reliability of the obtained results.

Finerenone is a novel, highly selective non-steroid MRA that exerts its action through binding to MR on cells (reviewed in ^68^). We observed typical, slightly rounded (or cobblestone-like) endothelial cell morphology in our control EoCs (Fig. 10 A). As expected, DKD mix (glucose 33 mmol/L, TNFɑ 10 ng/mL and aldosterone 0.1 nmol/L) induced changes in endothelial cell shape: HUVECs became more elongated (Fig. 10 B). Cells did not align in the direction of the applied flow. Changes in endothelial cell morphology indicate cytoskeleton reorganisation as a response to the stress, and, potentially, transition to an inflammatory phenotype.^69^

**Fig. 10.**
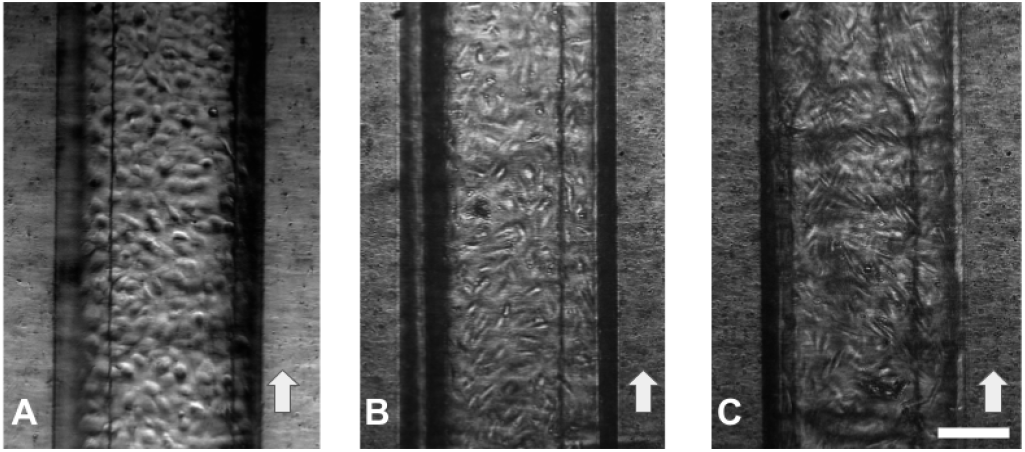
HUVEC morphology in an EoaC after 24 h of stimulation with DKD mix or DKD mix with finerenone (1 µM) under shear stress (10 dyn/cm^2^). A - typical cobblestone-like morphology can be observed in the control; B - endothelial cell morphology after 24 h of stimulation with DKD mix. Cell elongation can be observed. Cells were not aligned along the direction of the flow; C - endothelial cell morphology after 24 h of stimulation with DKD mix and finerenone (1 µM). Cell morphology is elongated. Cells are not aligned along the direction of the flow. The white arrows indicate the direction of the flow. Scale bar: 200 µm.

The addition of finerenone (1 µM) did not alter the DKD-induced endothelial cell elongation (Fig. 10 C). Cells maintained an elongated shape without aligning in the direction of flow.

#### Finerenone attenuates the increase in endothelial permeability in DKD environment with applied shear stress

We evaluated endothelial barrier permeability in an EoaC with applied shear stress of 10 dyn/cm^2^ after 24 h incubation in the DKD environment with or without adding finerenone (1 µM). Finerenone binds to the cell MRs. The MR has an important functional role in RAAS and is expressed in a variety of tissues, mostly in the kidneys and the heart. MRs are mainly activated by aldosterone and have an important effect on blood pressure, inflammation and fibrosis processes, oxidative stress and endothelial dysfunction (reviewed in ^70^). Aldosterone binding to the MR in the cytosol results in the transfer of this complex to the nucleus, targeting gene transcription (genomic actions of aldosterone) to increase the number of sodium channels on the apical side of the cells.^71^ Non-genomic MR-mediated effects of aldosterone include an increase in cytosolic calcium, production of ROS, and initiation of inflammatory pathways (reviewed in ^72,73^).

Elevated MR expression has been demonstrated in several pathological conditions, including hyperglycaemia and CKD. It has been shown that hyperglycaemia promotes MR glycosylation, thus augmenting its transcriptional activity.^74^ Elevated serum aldosterone levels have been reported in patients with CKD.^75^ Taken together, the overexpression of MR due to hyperglycaemia, in addition to elevated aldosterone levels, results in pronounced genomic and non-genomic effects on cells. Even though the research on the effects of aldosterone and MR activation has been focused mainly on tubular epithelial cells, mesangial cells and glomerular podocytes, the endothelial barrier layer is a crucial component of glomerular permeability barrier that is affected in DKD.

We observed a strong, statistically significant decrease in endothelial permeability of HUVECs in DKD milieu with the addition of finerenone compared to cells challenged with DKD mix only (mean P_app_ 8.64 × 10^-5^ ± 1.01 × 10^-5^ cm/s for DKD environment with finerenone in comparison to mean P_app_ 1.03 × 10^-4^ ± 0.1 × 10^-4^ cm/s for DKD environment only) (p = 0.0001 to 0.001) (Fig. 11).

**Fig. 11.**
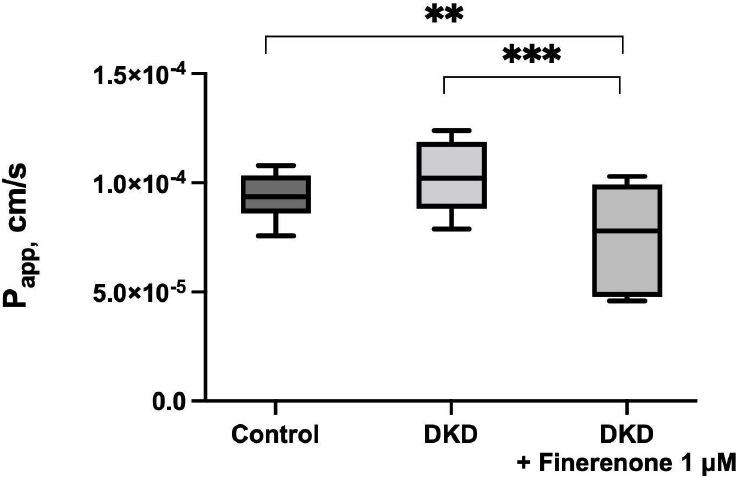
Endothelial permeability in DKD environment or DKD environment with added finerenone (1 µM) with applied shear stress (10 dyn/cm^2^) in an EoaC. The data represents two biological replicates each for control and DKD environment (n=2), and three biological replicates for DKD environment with finerenone (n=3). Seven images per biological replicate were obtained, and, as a result, 14 images were analysed for the control and DKD environment experiments, while 21 images were analysed for cells in a DKD environment with finerenone. Each box plot depicts a minimum and maximum value, as well as median and lower and upper quartiles of the data set. Level of significance indicated: ** - 0.001 to 0.01; *** - 0.0001 to 0.001.

The reduction in endothelial permeability due to MR blocking in DKD milieu is in agreement with previous studies considering the therapeutic effects of MR antagonists. Crompton and colleagues demonstrated a decrease in albuminuria in streptozotocin-induced diabetic rats after treatment with the MR antagonist spironolactone.^76^ This improvement in glomerular barrier function was observed along with decreased concentrations of the matrix metalloproteinases 2 and 9 (MMP2 and MMP9). As MMP2 and MMP9 have been shown previously to be the key glycocalyx sheddases,^41,77^ a restoration of the glycocalyx can be expected as a result of lowered MMP concentrations. The improvement in glomerular permeability barrier in this study could thus be linked to a restored glycocalyx. Moreover, the effects of MR antagonists on endothelial permeability are not limited to actions on endothelial glycocalyx only. Gonzalez-Blazquez and colleagues demonstrated >40% reduction in albuminuria in Munich Wistar Frömter (MWF) rats after treatment with finerenone. MWF rats are an experimental genetic model of spontaneous non-diabetic albuminuria and renal injury that mirrors features observed in patients with chronic kidney disease. These effects were achieved via enhancement of nitric oxide (NO) bioavailability and decreased superoxide anion levels (meaning reduced oxidative stress) after finerenone treatment.^78^ Another recent study by Gil-Ortega and colleagues demonstrated that both of these effects, the decrease in MMP2 and MMP9 activity and reduction of oxidative stress, take place in MWF rats, as well as reduced arterial stiffness, related to elastin reorganisation.^79^ These *in vivo* studies highlight the molecular mechanisms behind the beneficial effects of finerenone on endothelial cell health and barrier properties.

### Conclusions and future outlook

The main motivation driving OOC development is its potential for modelling a physiologically relevant microenvironment, thus ensuring more physiological cell response.^80^ This is crucial for elucidating the mechanisms of action in physiological and pathophysiological milieu. Better understanding of the diseases and the effects of pharmaceuticals enables the application of this knowledge in a personalised medicine. Moreover, accurate assessment of drug effects on target cells will speed up drug development and, eventually, will enable replacing the animal experiments.^81^

We have developed an EoaC, with a capacity to apply shear stress, that is suitable for endothelial permeability evaluation. We have demonstrated biocompatibility of the EoaC device with HUVECs, as well as good potential for long-term studies in this EoaC. We observed protective cell-ECM interactions between HUVECs and gelatin/mTG hydrogel in response to applied shear stress with regards to HUVEC morphology. The endothelial barrier permeability response to shear stress was consistent with the literature data. Further, we were able to mimic the DKD environment in the EoaC and observe changes in cell morphology, as well as barrier permeability as a response to DKD milieu. The obtained results on endothelial barrier layer response to DKD stimuli were in agreement with literature data. Moreover, a statistically significant improvement in endothelial permeability after addition on finerenone (1 µM) to DKD milieu allows us to conclude that this EoaC would be suitable for modelling the DKD environment *in vitro*, and holds a potential for use in evaluation of drug candidates targeted to endothelial layer resilience.

Further advancements in this EoaC should be considered. For instance, the heterogeneity of the endothelium throughout the body is well recognized.^82-84^ Differences in the local environment depending on the organ and the area of the vascular bed results in phenotypic differences and even some variability of the cell responses. Human glomerular endothelial cells (hGEnCs) could be considered in an optimised EoaC for studying DKD, to avoid cell responses potentially influenced by endothelial heterogeneity. Moreover, patient-derived hGEnCs in an EoaC would enable evaluation of patient-specific responses and tailoring the most effective medication regimen to address albuminuria. The use of induced pluripotent stem cell-derived endothelial cells (iPSC-EC) would allow the development of a more personalised EoaC approach.^85,86^ Regarding experimental conditions, several aspects could be considered. For instance, longer experiments would allow better characterisation of the endothelium in a chronic pathophysiological environment, such as diabetes and DKD. Mimicking natural fluctuations of the glucose levels over the day would provide a further refinement of this test system. Moreover, this versatile EoaC could be applied for modelling a variety of pathophysiological conditions affecting endothelial barrier function, for instance, sepsis, pulmonary edema, and others.

