## Supporting Information for "Endothelium-on-a-chip for permeability studies in diabetic and diabetic kidney disease milieu"

### Preparing the flow microchannels in PDMS

*Designing the microchannel and preparation of SU-8 50 master mould*

The microchannel design was prepared using CleWin software (WieWeb, The Netherlands). The pattern of the microfluidic channel was printed as a negative mask (ProArt B.V., The Netherlands). This pattern was transferred onto a negative photoresist layer deposited on a Borofloat glass wafer (Borofloat 33, diameter 100 mm; Handelsagentur Helmut Teller, Germany).

To prepare 100-µm-high structures, SU-8 50 photoresist (Kayaku Advanced Materials, Inc., Germany) was spin-coated on a wafer to the desired thickness using a CEE™ spin coater (Brewer Science, MO, USA). A soft bake step was performed with the spin-coated wafer on a hotplate with the following programme:

1) heating from 20°C to 65°C in 45 min (1°C/min);
2) incubating at 65°C for 10 min;
3) heating from 65°C to 95°C in 30 min (1°C/min);
4) incubating at 95°C for 30 min.

After removing the wafer from the hotplate and passively cooling down to room temperature, the mask was placed on top of the wafer and was exposed to the UV light (365 nm, 250 mJ/cm$^2$) (OAI exposure system, CA, USA) for 21.9 sec. A post-exposure bake was then performed on the hotplate with the following programme:

1) heating from 20°C to 65°C in 45 min (1°C/min);
2) incubating at 65°C for 1 min;
3) heating from 65°C to 95°C in 30 min (1°C/min);
4) incubating at 95°C for 10 min.

After passively cooling down to room temperature as described before, the wafer was inserted in a glass Petri dish with SU-8 developer (Kayaku Advanced Materials, Inc., Germany) for 15 min to wash away the unexposed SU-8 50. An extra baking step was then performed by placing the wafer with the structures up on a hotplate and incubating it at 150°C for 20 min, followed by cooling it down passively to 65°C. This step repairs potential cracks in the structures, as well as makes them stronger.[1]

To facilitate easier removal of PDMS from the master mould, as well as extend its lifetime, a silanisation step was performed. To silanise the mould, the mould wafer with structures was placed under vacuum for 1 h together with a 10 µL of perfluorooctyltrichlorosilane (PFOCTS) (Sigma-Aldich, Germany) dropped on a glass slide. After silanisation the mould surface became hydrophobic, facilitating easier peeling of the PDMS from the master mould.

*Casting the microchannels in PDMS*

The master mould was fixed in a rectangular Petri dish with a piece of tape, and 70 g of a degassed solution of PDMS monomer and curing agent (10:1) was poured onto it. This produced the top PDMS layer of the EoaC device at a final thickness of about 5 mm. The Petri dish was placed on a hotplate to cure the PDMS at 70°C for 1.5 h. Afterwards, the top PDMS layer was cut with a scalpel to a size of about 4.5 x 2 cm. Inlet and outlet holes were punched with a 1-mm biopsy puncher. To protect it from environmental dust, each microfluidic channel was sealed with Scotch tape until the EoaC device layers were bonded together.

#### Gelatin/mTG hydrogel preparation in the hydrogel channel

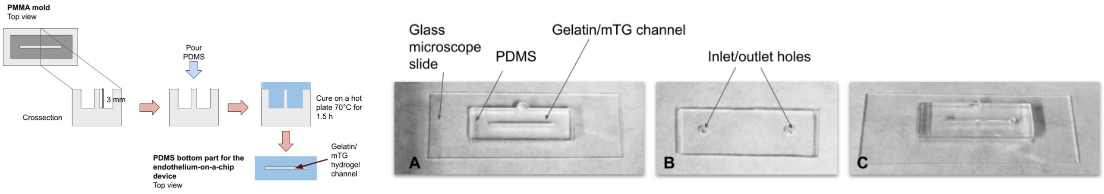


***Figure S 1*** *Preparation of the gelatin/mTG hydrogel channel in the lower PDMS layer of the EoaC device. Left side: PDMS elastomer was mixed with a curing agent (10:1) and poured in the PMMA mould. It was then cured on a hot plate at 70°C for 1.5 h. After curing, the PDMS part was manually peeled from the mould. This way the lower PDMS layer with a channel for the hydrogel for the EoaC device was obtained. Right side: A - the lower PDMS layer with the channel for the gelatin/mTG hydrogel is bonded to a microscope slide to mechanically support the hydrogel channel. This will later prevent gelatin/mTG hydrogel from accidental damage; B – the PDMS cover sheet with punched inlet/outlet holes (diameter: 2 mm); C – the PDMS cover sheet positioned on the lower PDMS layer and aligned with the gelatin/mTG hydrogel channel. The device is now ready to be filled with the gelatin/mTG solution to form the hydrogel.*

#### Cell maintenance and seeding in the EoaC device

Commercially available human umbilical vein endothelial cells (HUVEC) (Lonza, Switzerland) from passage 5 to 8 were used. HUVECs were maintained in endothelial cell culture medium in 37°C and 5% $CO_2$ in T75 flasks (Corning, AZ, USA) with 0.1 % gelatin coating. Endothelial cell medium was changed every two to three days. Gelatin coating was prepared prior to cell seeding in the flask by diluting 2% gelatin solution (Type B) (Sigma-Aldrich, Germany) with sterile 1x phosphate-buffered saline (PBS, ThermoFisher Scientific, NY, USA) to 0.1% gelatin and introducing 2 mL of this solution in a T75 flask. The flask with 0.1% gelatin solution was incubated at 37°C for 45 min, after which excess gelatin solution was removed by aspiration. The flask was then incubated at 37°C for 30 min to allow the flask surface to dry before introducing the HUVECs. The gelatin coating of the polystyrene cell growth surface in the flask mimics the presence of ECM proteins and improves cell attachment and survival.

For cell seeding in the EoaC devices, HUVECs were detached from the growth surface by adding 2 mL of 0.25% trypsin/ethylenediamine tetraacetic acid (EDTA) solution (ThermoFisher Scientific, NY, USA) per T75 flask and incubating at 37°C in 5% $CO_2$ for 5 min. After cell detachment, the activity of trypsin/EDTA was inhibited by the addition of 4 mL of 10% foetal bovine serum (FBS) in 1x PBS. The cells were separated from the suspension by centrifugation at 250 rcf for 5 min at RT and subsequent removal of the supernatant. The cell pellet was resuspended in cell growth medium and cells were counted using a Neubauer chamber (OPTIK-Labor, Germany). Cell suspension was diluted with endothelial cell culture medium to obtain concentration of 5 x $10^6$ cells/mL for cell seeding in an EoaC device.

The cell suspension was injected gently and slowly into the inlet of the EoaC device using a 1-mL syringe without needle. The EoaC devices were left undisturbed for 15 min at RT in the laminar flow hood to allow HUVECs to settle on top of the gelatin/mTG hydrogel. The EoaC was then placed in a cell incubator at 37°C and 5% $CO_2$ to allow the endothelial cells to attach to the surface of the gelatin/mTG hydrogel and form a monolayer.

Two hours after cell seeding, the EoaC was inspected under the microscope to evaluate how many cells had attached to the hydrogel. Fresh endothelial cell culture medium was gently introduced with a 1-mL syringe without needle into the microchannels to rinse out any unattached cells, and to supply the attached cells with fresh nutrients. This encouraged the formation of an endothelial cell monolayer. A 100-μL pipette tip was inserted in the outlet to contain and remove the unattached cells. About three hours after cell seeding, a quality control of the EoaC was performed by evaluating the confluency of cells under the microscope. The confluency was evaluated visually, by monitoring the coverage of the endothelial cell monolayer on the growth surface in the cell culture chamber. Only EoaC that had a confluency of 95% or higher were used in permeability assays.

**Flow system setup: components and application of the flow to the EoaC**

Sterile syringes (volume 60 mL) (BBraun, Germany) were secured in the syringe pumps (Prosense B.V., The Netherlands). Syringes were connected to the inlet tubes with connectors (Becton Dickinson, NJ, USA). The connectors allowed the exchange of syringes more easily without introducing air bubbles in the system. Inlet and outlet tubes were prepared from a Teflon tube (0.8 x 1.6 mm, inner and outer diameter) (Polyfluor Plastics B. V., The Netherlands). To secure the inlet tube in the connector, a syringe needle (outer diameter 0.8 mm) with a Luer Lock (BBraun, Germany) was cut to about 1 cm in length. The tip of the cut needle was then deburred using sandpaper and carefully inserted into the inlet tube. The Luer Lock then allowed the inlet tube to be secured in the connector.

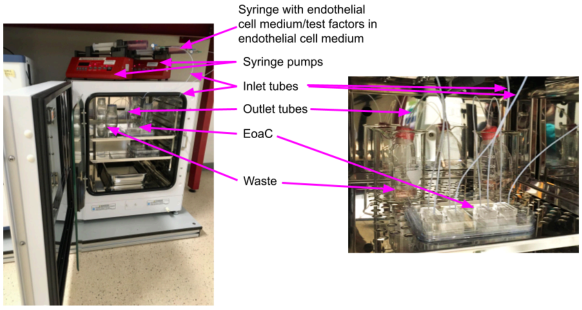


***Figure S 2** The flow system setup for evaluating endothelial permeability in DKD milieu in the EoaC. The flow system setup consists of syringe pumps, syringes filled with cell medium or test factors in cell medium, inlet and outlet tubes, as well as EoaC, and waste containers.*

To prepare the inlet and outlet tubes for connecting to the EoaC inlet and outlet, a syringe needle (about 1 cm long, outer diameter 0.8 mm) was prepared as described above and carefully inserted into the inlet and outlet tubes. The needle was inserted halfway into the tube. That way, the tubes can be connected to an EoaC by inserting the other end of the needle into an inlet or outlet hole of the device. PDMS is an elastic material and closely surrounds the needle tip, which ensures leak-free connections.

Inlet and outlet EoaC tubes were sterilised with 70% ethanol at RT for 30 min and rinsed thoroughly with sterile 1x PBS three times before and after every use. Sterilised tubes were stored in a sterile plastic box between the experiments.

#### Imaging and result analysis

Seven fluorescence microscopy images of each EoaC along the length of the cell cultivation chamber were taken (Fig. S 3 A and B). The focus was set on the surface of the gelatin/mTG hydrogel. Additionally, brightfield images were taken to evaluate endothelial cell morphology.

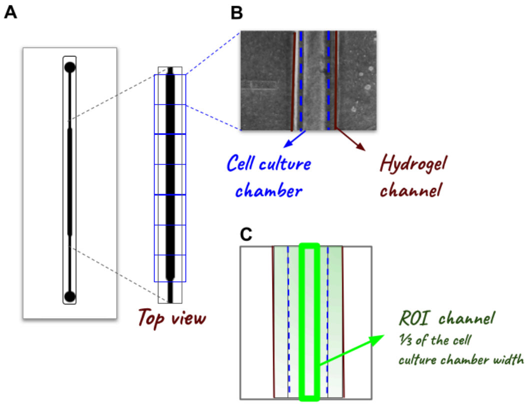


***Figure S 3*** *A - the pattern for taking fluorescence images of the EoaC for fluorescence-based permeability assay. Seven images were taken for each EoaC. The total imaged area covered the length of the cell culture chamber; B - the microphotograph of an EoaC device depicting the hydrogel channel and cell culture chamber alignment; C - ROI is set as ⅓ of the cell culturing chamber width and manually positioned in the middle of the cell culture chamber.*

The green channel of the fluorescence microscopy images was used for fluorescence intensity analysis with the ImageJ program (version 1.53a, NIH and University of Wisconsin, WI, USA). A region of interest (ROI) of 1/3 of the width of the cell culture chamber was set (w=94 px, h=1040 px) (Fig. 4 C). This region was manually positioned in the middle of the cell culture chamber. Intensity density (IntDen) measurements were obtained for each ROI in arbitrary units (a.u.). To correct for the background fluorescence, the background fluorescence intensity measurements were obtained in each image using an ROI having the same dimensions as above, but then positioned outside of the channel area. Fluorescence intensity for each image was corrected by subtracting the measured background fluorescence intensity from the corresponding fluorescence intensity in the cell culture channel.

#### Live/dead well-plate assay with atrasentan and finerenone

To select target concentrations of atrasentan (Bio-Connect, The Netherlands) and finerenone (Bio-Connect, The Netherlands) for our EoaC experiments, we first performed a live/dead assay on HUVECs in a well plate. The initial concentrations of atrasentan (0.5 µM) and finerenone (1 nM and 10 nM) were selected based on the literature data. For the live/dead assay we used calcein AM (excitation max 494 nm, emission max 517 nm) (ThermoFisher Scientific, NY, USA) in combination with propidium iodide (PI) (excitation max 535 nm, emission max 617 nm) (ThermoFisher Scientific, NY, USA).

For the live/dead assay, gelatin hydrogel crosslinked with microbial transglutaminase (gelatin/mTG hydrogel) was prepared in a 24-well plate (Greiner Bio-One, Germany) according to a previously described method (see in Experimental: *Gelatin/mTG hydrogel* *preparation in the hydrogel channel*) with the only difference being that the gelatin/mTG solution was filled into a 24-well plate, at a volume of 0.5 mL/well. After gelatin/mTG crosslinking, sterilisation and conditioning steps, HUVECs at a concentration of 150 000 cells/mL in endothelial cell medium were seeded on gelatin/mTG hydrogel, at a volume of 1 mL per well. Cells were placed at 37°C in 5% $CO_2$ to allow cell adherence and monolayer formation.

After about 24 hours, a DKD mix (glucose: 33 mmol/L; aldosterone: 0.1 nmol/L; and TNFα 10 ng/mL; in the endothelial cell medium) with or without various concentrations of atrasentan and finerenone was added to HUVECs (see Table S 1). The stock solutions of atrasentan and finerenone were prepared by diluting the pharmaceuticals in dimethyl sulphoxide (DMSO) to a concentration of 0.5 mg/mL. The test concentrations of pharmaceuticals were prepared by diluting the stock solution with endothelial cell culturing medium to the desired concentration. We prepared an additional control for the highest concentrations of atrasentan (50 µM and 500 µM) to prevent the effects of high dilution of DKD mix with atrasentan stock solution. These controls were prepared by substituting the according volume of atrasentan stock solution in the DKD mix with the endothelial cell medium. High dilution of the DKD mix could result in altered results and incomplete conclusions about the effects of atrasentan on cell viability.

| Pharmaceutical substance | Concentrations |
|---|---|
| Atrasentan | 0.05 µM; 0.5 µM; 5 µM; 50 µM; 500 µM |
| Finerenone | 1 nM; 10 nM; 100 nM; 1 µM; 10 µM |

***Table S 1*** *Concentrations of pharmaceuticals for live/dead assay.*

After 24 h of stimulation, HUVECs were rinsed with sterile 1 x PBS. Then, 1 mL of calcein AM (2 µg/mL) and PI (2 µg/mL) solution in 1x PBS was added to each well and incubated at room temperature (RT) for 10 min. Wells were rinsed three times with 1x PBS. Cell culturing medium (0.5 mL/well) was added to the wells. Cells were visualised with a Leica DMIL LED microscope, light source EL6000, and camera Leica DFC310 FX (Leica, Germany). Fluorescence images were taken. One biological replicate was obtained for the live/dead assay for each concentration of pharmaceuticals, and three technical replicates (wells) were tested per biological replicate. Then, one image from each well was selected for further analysis (n=3). To determine the % of apoptotic cells, live and apoptotic cells were counted using the ImageJ program (version 1.53a, NIH and University of Wisconsin, WI, USA), and the percent of apoptotic cells was calculated.

Diabetic and DKD environments *in vivo* are a complex mix of endogenous factors. For this reason, we merged all DKD factors studied in our experiments in a DKD mix (glucose 33 mmol/L, TNFα 10 ng/mL, aldosterone 0.1 nmol/L). We then compared whether addition of the pharmaceuticals in different concentrations decreased the percentage of apoptotic cells in comparison to cells exposed to DKD mix only.

We initially selected three concentrations of atrasentan and finerenone based on the data in literature. Boels and colleagues have demonstrated the protective effect of 0.5 µmol/L atrasentan on the glycocalyx thickness of HUVECs co-cultured with human brain pericytes under 10 dyn/$cm^2$ shear stress after stimulating them with diabetic patient serum for three days.[2] Dutzmann and colleagues tested the finerenone (1 nM, 10 nM) in combination with aldosterone and observed that finerenone reversed aldosterone-induced HUVEC apoptosis in a dose-dependent manner.[3] Based on this, we decided to test the effect of 0.05 μM, 0.5 μM and 5 μM atrasentan and 1 nM, 10 nM and 100 nM finerenone in combination with the DKD environment.

We observed 2.58 ± 0.35% apoptotic cells in control in comparison to 13.57 ± 4.60 % apoptotic cells in the DKD environment (Fig. S 2 C). This result was statistically significant ($p < 0.05$). However, the addition of atrasentan and finerenone in selected concentrations to the DKD mix did not reduce endothelial cell apoptosis.

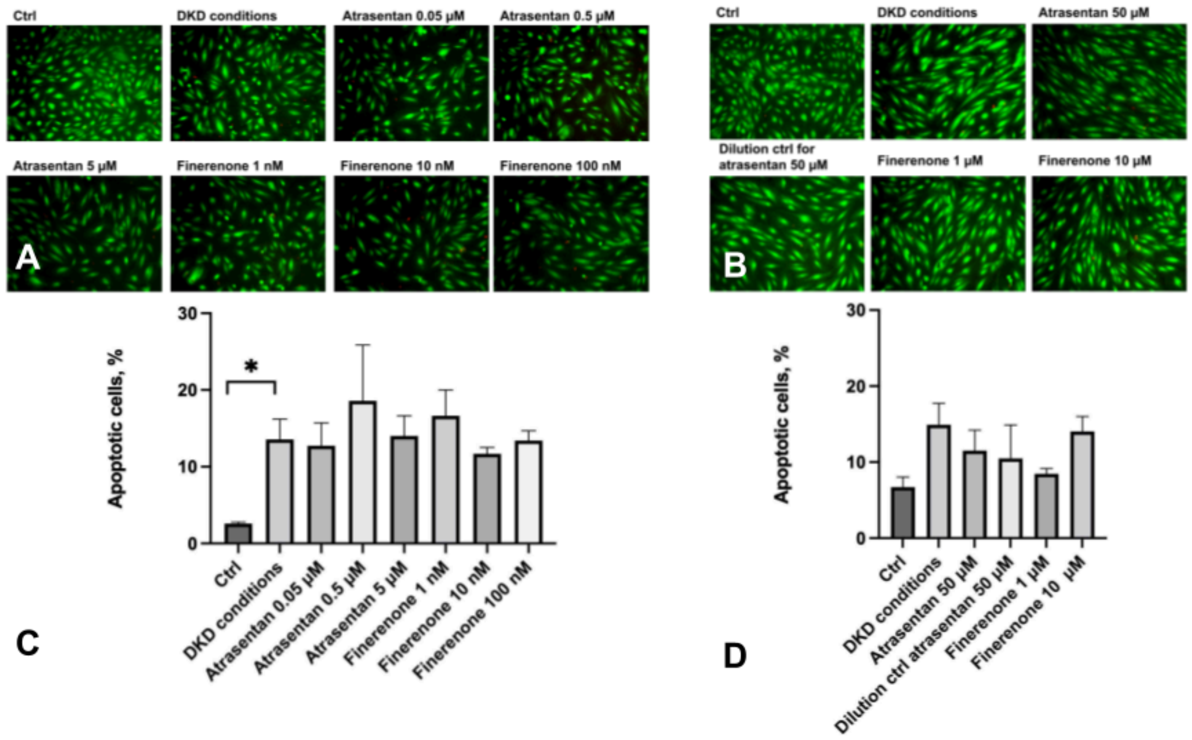


***Figure S 2*** *Live/dead assay on HUVECs in a static system in DKD environment (using a mix of DKD factors) and in DKD environment with added pharmaceuticals atrasentan and finerenone in several concentrations. A - representative fluorescence microscopy images of the live/dead assay with the DKD mix and DKD mix with atrasentan (0.05 μM; 0.5 μM; 5 μM) or finerenone (1 nM; 10 nM; 100 nM); B - representative fluorescence microscopy images of the live/dead assay with DKD mix and DKD mix with atrasentan (50 μM; dilution ctrl for 50 μM) or finerenone (1 μM, 10 μM); C - comparison of apoptotic cells (%) in control, DKD environment or in DKD environment with added atrasentan (0.05 μM; 0.5 μM; 5 μM) or finerenone (1 nM; 10 nM; 100 nM); D - comparison of apoptotic cells (%) in control, DKD environment or in DKD environment with added atrasentan (50 μM; vehicle for 50 μM) or finerenone (1 μM, 10 μM). The result was obtained from one biological replicate with three technical replicates (n=3). Green - calcein AM, red - PI. Level of statistical significance indicated: * - <0.05.*

We then decided to increase the concentrations of atrasentan and finerenone. Accordingly, for our second experiment we compared apoptotic cells in the DKD environment vs. DKD environment with 50 μM and 500 μM atrasentan along with dilution controls, or 1 μM and 10 μM finerenone. We observed a cytotoxic effect of 500 μM atrasentan since all of the HUVECs detached from the growth surface (result not shown). We observed 6.70 ± 2.31 % apoptotic cells in control in comparison to 14.92 ± 4.88 % apoptotic cells in the DKD environment (Fig. S 1 D). We did not observe reduced endothelial cell apoptosis after addition of atrasentan. However, a decrease in cell apoptosis (8.46 ± 1.21 %) was observed after addition of 1 μM finerenone to a DKD mix, compared to the DKD environment without an added pharmaceutical. However, this effect did not reach statistical significance.

Based on our results from the live/dead assay, we decided to proceed with 1 μM finerenone in DKD mix for our further EoaC studies.
